# Examining Relationships between Functional and Structural Brain Network Architecture, Age, and Attention Skills in Early Childhood

**DOI:** 10.1101/2024.10.03.616504

**Authors:** Leanne Rokos, Signe L. Bray, Josh Neudorf, Alexandria D. Samson, Kelly Shen, Anthony R. McIntosh

## Abstract

Early childhood is a critical period showing experience-dependent changes in brain structure and function. The complex link between the structural connectivity (SC) and functional connectivity (FC) of the brain is of particular interest, however, its relationship with both age and attention in early childhood is not well understood. In this study, children between the ages of 4 and 7, and at a one-year follow-up visit, underwent neuroimaging (diffusion-weighted and passive-viewing functional magnetic resonance imaging) and assessments for selective, sustained, and executive attention. We examined regional graph theory metrics and SC-FC coupling of the structural and functional networks. Partial least squares (PLS) was used to investigate longitudinal brain measure changes and cross-sectional associations with age and attention. We observed longitudinal changes in functional graph theory metrics and age-related decreases in SC modularity. Region-wise graph theory analyses revealed variable brain-behaviour relationships across the brain, highlighting regions where structural topology is linked to age and attentional performance. Furthermore, we identified SC as a dominant predictor of age when compared to FC and SC-FC coupling. The findings emphasise how early childhood is a dynamic period where cognitive functioning is intricately and predominantly linked to structural network features.

**Significance Statement:** This study investigates early childhood brain development, particularly the changes in structural and functional connectivity of the brain and the relationship between them. We examined children aged 4 to 7 over a year, and used graph theory analyses to characterise variable developmental changes and brain-behaviour relationships across the brain. Our findings emphasise the nuanced and regionally specific relationship between brain network structure and behaviour in early childhood, highlighting regions where structural topology correlates with attentional performance. This research underscores the dynamic nature of childhood brain development and the predominant role of SC in cognitive maturation, providing valuable insights into typical developmental trajectories and potential targets for early intervention strategies.

## 1. Introduction

Early childhood is a critical period of cognitive, behavioural, and brain development (Brown and Jernigan, 2012). Executive functioning skills, including working memory and attention, undergo intertwined development as children experience increasing and new attentional demands. Top-down attention skills have been classified into three attentional sub-systems that demonstrate unique developmental changes (Zhan et al., 2010). These processes include sustained attention (i.e., *maintaining* attention for extended durations), selective attention (i.e., *orienting* attention for locating a target among distractors), and executive attention (i.e., *controlling* or shifting responses to changing circumstances; Breckenridge et al., 2013; Manly et al., 2001). Characterising brain developmental changes underlying each attentional process has implications for understanding healthy and atypical development. A cross-sectional study with typically developing girls reported a link between distinct intrinsic functional connectivity (FC) networks and specific attentional abilities, as well as age-related FC network differences (Rohr et al., 2018). Across early childhood, there are also white matter (WM) changes, including alterations in mean, radial and axial diffusivity, fractional anisotropy, and more minor changes in streamline counts (i.e., structural connectivity measures; Dimond et al., 2020; Lim et al., 2015). Diffusion-weighted magnetic resonance imaging (offering an indirect measure of WM microstructure) has been used to characterise non-linear, regionally heterogeneous changes in WM tracts across the lifespan (Lebel et al., 2019, 2012; Reynolds et al., 2019). A longitudinal approach for assessing both functional and structural topology of early childhood brain networks will provide a more complete understanding of these attentional processes in typical development.

Brain network topology assessed with graph theory treats regions as nodes and structural and functional connections as edges (Bullmore and Sporns, 2009; Rubinov and Sporns, 2010). The brain appears to display a “small-world” topology, balancing segregation and integration for efficient information processing with high clustering (i.e., regions tightly connected with their neighbours) and short paths (Watts and Strogatz, 1998). The brain’s community structure (i.e., nodal organisation into interconnected groups) can also be assessed with a modularity measure quantifying how well a network can be separated into clear, non-overlapping modules (Newman, 2004). Clustering and modularity metrics have been used to characterise developmental topology in numerous studies (Cao et al., 2017; Fair et al., 2009; Hagmann et al., 2010; Supekar et al., 2009; Wierenga et al., 2016). Long-range axonal projection maturation is believed to underlie developmental increases in structural integration (e.g., more long-distance and intermodule connections) and decreasing segregation (e.g., fewer short-distance and intra-module connections; Cao et al., 2017; Hagmann et al., 2010; Tymofiyeva et al., 2013; Vohryzek et al., 2020). Studies have also found diverse functional connectivity profiles, including changes in functional segregation and integration with age (Fair et al., 2009; Marek et al., 2015; Pines et al., 2022; Tooley et al., 2022). The variable patterns of segregated and integrated functional networks may depend on the network’s position on the sensorimotor-association axis (Keller et al., 2023), highlighting the importance of considering region-wise metrics when investigating the relationship between network development and attentional skills.

The relationship between structural and functional connectivity is also of immense interest (Suárez et al., 2020 for review). Studies have suggested that the relationship between SC and FC strengthens across the lifespan (Betzel et al., 2014; Hagmann et al., 2010). SC-FC coupling has also been found to be a stronger predictor of age (between 18-82 years) compared to SC or FC alone (Zimmermann et al., 2016). Conversely, there are reports that SC did not have a large impact on the decline in FC with ageing (Madden et al., 2020; Tsang et al., 2017). Recent work has further suggested nonlinear, heterogeneous structure-function relationships across the brain (Baum et al., 2020; Vázquez-Rodríguez et al., 2019; Zamani Esfahlani et al., 2022). Therefore, studying both SC and FC changes, including region-wise metrics, can help elucidate healthy brain development patterns along with potential predictive factors of behavioural outcomes.

In this study, we investigated structural and functional networks of children between the ages of 4 and 7, and at a one-year follow-up visit. We used partial least squares (PLS) to assess if regional graph theory metrics and SC-FC coupling changed over the follow-up period (longitudinal), and to explore if the brain measures were related to age and attention (cross-sectional behaviour/age associations at two time points). In a combined analysis, we compared SC, FC, and SC-FC coupling with age. We hypothesised that older age and better attention skills would be associated with lower, region-specific segregation and greater SC-FC coupling.

## 2. Materials and Methods

### 2.1 Participants

Data were collected from typically developing children (4-8 years) at Alberta Children’s Hospital. All participants provided written informed consent (parents) and assent (children). The study received approval by the Conjoint Health Research Ethics Board of the University of Calgary. Exclusion criteria for potential participants included: history of neurodevelopmental or psychiatric disorders; neurological diagnosis; chronic medical condition or gestational age less than 37 weeks. The inclusion criteria were participants who had three neuroimaging scans (i.e., T1 weighted, multishell dMRI, and fMRI scans). Following neuroimaging processing and quality control exclusion criteria described below, the final sample included 39 participants (20 girls) with an initial scan (Mean age=5.76 [4.14-6.88]) and a scan at one-year follow-up (Mean age=6.85 [5.13-7.89]) (i.e. 78 scans). Four participants were reported to be non-right-handed by their parents including left-handed (*n*=1), more left-handed at time point one and left-handed at follow-up (*n*=1), and ambidextrous (*n*=2) at time point one but more right-handed at follow-up.

### 2.2 Data Collection

During two distinct sessions (2-hours) within a two week period, both cognitive assessments and MRI scans were conducted. The initial session consisted of a number of attention tasks that were administered in a randomised order (within and between sessions). The participants also underwent training in an MRI simulator whereby they practised lying down in the scanner while listening to MRI scanner sounds through headphones. During the practice scan they also viewed the same 18-minute video shown in the real MRI scanner, such that children had similar familiarity with the video at the baseline and follow-up scans. The actual MRI scanning process and the remaining attention tasks were conducted during the second session. The attention tasks were completed in a testing room adjacent to the MR simulator.

### 2.3. Assessment of Attention Skills

Participants’ attention skills were assessed with four tasks including measures of sustained attention (visual and auditory), selective attention, and executive attention. The assessments were modelled on the Early Childhood Attention Battery (Breckenridge et al., 2013), which was developed as an adaptation of the Test of Everyday Attention for Children (TEA-Ch; Manly et al., 2001) suitable for children aged 3 to 6 years old. All assessments were administered using a Dell laptop computer (screen size=31cmx17.5cm; viewing distance=35-50cm), except for the selective attention subtest (see Section 2.3.2). External speakers were used to play auditory items. To ensure the child comprehended the instructions for each computer-based task, a practice trial was completed, and repeated if necessary.

#### 2.3.1. Sustained Attention Tasks

The visual sustained attention task included the presentation of a continuous sequence of pictures, with each image appearing for 200ms (inter-stimulus interval; ISI=1800ms). Thirty target images (an animal) and 120 non-targets (common everyday items) were presented. When a target appeared, the child’s task was to respond “yes”, “animal”, or the name of the animal. If four consecutive targets were missed, the child received a prompt to maintain their attention. The measure was scored as the sum of correct responses, minus errors and prompts.

The auditory sustained attention task included a continuous sequence of words (average duration= 650 ms, ISI=1350 ms). Both mono-syllabic target (animal) and non-target (familiar item) words were presented. The child’s task was to respond “yes”, “animal”, or the name of the animal when they heard a target word. The number of correctly identified targets, minus errors and prompts, was taken as the auditory score. Lastly, the total sustained attention score was calculated as the mean of visual and auditory sustained attention scores.

#### 2.3.2 Selective Attention Task

In the selective attention task, children were presented with a laminated letter-sized search sheet containing both targets (18 red apples) and distractors (162 white apples and red strawberries). They were given 60s to point to the targets and an experimenter marked correctly identified targets with an erasable marker. The score was the sum of correctly identified targets.

#### 2.3.3 Executive Attention Task

A child-appropriate Wisconsin Card Sorting test adaptation was used for the executive attention task (Robinson et al., 1980). The child’s task was to determine which type of balloon a teddy bear preferred. The bear liked a specific colour of balloon in stage one and a different colour in stage two. In stage three, the bear liked a specific shape of balloon. Visual feedback (and no other information) was given to the children on whether they made the correct choice. Six consecutive correct responses (out of 20 possible trials for each stage) were required to pass. The test was discontinued if a child failed a stage. The executive attention task score was the total number of incorrect or incomplete trials (multiplied by negative 1 so positive values reflected better attention for all analyses).

#### 2.3.4 Behavioural Analyses

The statistical analyses were conducted using Matlab Version 9.11 (R2021b) and R version 4.3.0 (R Core Team, 2023). The relationships between age (fixed effect) and attention skills were computed with the “lme4” package in R (Bates et al., 2015) to account for repeated measures (i.e., participants’ intercepts specified as random effects). The Satterthwaite method (“lmerModLmerTest”) was used to calculate the *p*-values for these models (Hrong-Tai Fai and Cornelius, 2007; Kuznetsova et al., 2017).

Multiple paired t-tests were conducted to explore whether there were differences in the performance of each attention assessment between time points. For a single participant that had a sustained attention score greater than 3 standard deviations from the mean, a single attention assessment (auditory) was used to calculate the average score. Additionally, one participant did not complete the executive attention task and thus the missing value was replaced with the sample’s average score.

### 2.4 Neuroimaging Data & Pre-processing

Participants underwent an MRI scan while awake and watching clips from a children’s television show (Elmo’s World). The scans were collected on a 3T General Electric MR750w scanner (GE, Waukesha, WI), with a 32-channel head coil. FMRI data were acquired with a gradient echo planar imaging pulse sequence (TR/TE=2500/30 ms; FA=70; FOV=64×64 mm; number of slices=34; resolution=3.5 mm isotropic voxels; number of volumes=437). T1-weighted imaging data were acquired with a FSPGR BRAVO sequence (TR/TE=6.764/2.908 ms; FA (flip angle)=10; FOV=512×512 mm; number of slices=226; resolution=0.8×0.8×0.8 mm isotropic voxels; scan time=577 s). Diffusion-weighted images were acquired using a 2D spin-echo EPI sequence (TR/TE=1000/86 ms; FA=90; FOV=230×230 mm; resolution=2.5 mm isotropic voxels; slices=45; 3 interspersed b=0s/mm^2^ volumes; b-values=1000, 2000s/mm^2^, number of directions=45; distributed on a whole-sphere).

The MRI data were preprocessed with TheVirtualBrain-UK Biobank pipeline (Frazier-Logue et al., 2022), which was adapted for paediatric populations. This multi-modal (anatomical, fMRI, dMRI) processing pipeline primarily relies on the FMRIB (Functional MRI of the Brain) Software Library toolbox (Jenkinson et al., 2012) to generate the structural and functional connectivity matrices used in subsequent analyses. The pipeline also produces detailed quality control (QC) reports of raw, intermediate and processed outputs of the pipeline, supporting the extensive manual QC of paediatric data. The pipeline steps are described below.

#### 2.4.1 Structural Processing Sub-pipeline

Preprocessing included an initial brain extraction with optiBET, an optimised brain extraction tool that results in high-quality, robust brain extraction for brains with severe pathology (Lutkenhoff et al., 2014). Processing also included a nonlinear registration of the T1-weighted images to an age-specific template (Fonov et al., 2011), NIHPD asymmetrical (natural paediatric template optimised for ages 4.5–8.5 years), followed by bias correction and segmentation. A 200 cortical regions-of-interest (ROI) parcellation (Schaefer et al., 2018) was transformed into the NIHPD template space and was registered to the T1-weighted images. Segmentation of white matter (WM), grey matter (GM) and cerebrospinal fluid signals was completed using FSL’s FAST. The WM and GM segmentations were used to create WM interface masks (i.e. the WM voxels bounding the GM) and exclusion masks.

#### 2.4.2 Functional MRI Processing Sub-pipeline

For the fMRI data, brain extraction, motion correction (MCFLIRT), slice timing correction, spatial smoothing (FWHM: 4 mm), high-pass filtering (100s), and registration (to the T1-weighted image and age-specific template) were completed using FSL’s FEAT toolbox. Manual classification of noise and signal components from FSL’s MELODIC ICA was completed and validated by two other lab members (KS, ADS) to create a training set of 23 participants that were representative of the dataset (e.g., matched for sex, time point) (Griffanti et al., 2017). The training set included 13 female and 10 male participants (Mage= 6 years [4.26-7.89], SD=1.00). FSL’s FMRIB’s ICA-based Xnoiseifier (FIX) with “aggressive” artefact removal (Griffanti et al., 2014) was used to perform denoising (i.e., all noise components and motion confounds were regressed). The aggressive approach removes the total shared variance between signal and noise and was adopted due to the presence of high levels of motion. The ROI parcellation registered to the T1-weighted images (i.e. as described above and output from the structural sub-pipeline) was registered to a reference fMRI volume.

#### 2.4.3 Diffusion MRI Processing Sub-pipeline

The diffusion data processing entailed B0 field estimation and unwarping using the Synb0-DisCo tool (Schilling et al., 2019) to help improve T1-weighted and dMRI image registrations. Specifically, the Synb0-DisCo tool created a synthetic undistorted B0 image for dMRI distortion correction (i.e., input into FSL’s TOPUP toolbox). Head motion and eddy correction (EDDY), brain extraction (BET), and registration of the templates, interface, and exclusion masks to DTI space were also completed. Diffusion tensor fitting (DTIFIT) with the FMRIB’s Diffusion Toolbox (FDT) was used to calculate the diffusion tensor model of each voxel, to check the fibre directions.

FDT’s BEDPOSTX was also implemented to model the crossing of fibres in each voxel and distributions of diffusion parameters by using Bayesian estimation (i.e., Markov chain Monte Carlo sampling). Probabilistic tractography using PROBTRACKX2 (fibre volume threshold=0.01; number of steps=2000; step length=0.5 mm; number of samples=5000; loop check=on) was then completed.

Specifically, PROBTRACKX2 takes many samples from the voxel-wise diffusion parameter distributions to create a histogram of the number of streamlines connecting each ROI pair. The interface masks (created previously as described in Section 2.4.1) were used to prevent over-defining white matter structures. The ROI masks were used as seeds while the GM exclusion masks were used to exclude any streamline that entered the masked region. For a particular ROI, their streamlines terminated when they reached any of the other ROI masks.

#### 2.4.4 Connectivity Matrices

The functional connectivity (FC) matrices were created for each participant by calculating the Pearson’s correlation coefficient of the BOLD time series for each pair of regions. The FC matrices were then Fisher Z-transformed. The SC weights matrix for each participant consisted of the number of streamlines between ROI pairs divided by the total number of streamlines seeded in a particular ROI. The matrices were symmetrized as the tractography approach does not provide information about connection directionality. The raw dMRI images’ FOV were consistently missing 7 ROIs due to regions being cut off at the bottom of the image stack, therefore those regions were excluded and the resulting 193×193 ROI functional and structural connectivity matrices were used for network analyses. No ROIs were excluded due to fMRI data quality. Consensus thresholding was applied to the SC matrices to address false positives that can result from probabilistic tractography (Shen et al., 2019). Specifically, connections were set to zero if they were not present in at least 75% of the high-quality dMRI scans (i.e., scans that passed the quality control steps described below; *n*=176) (Roberts et al., 2017).

#### 2.4.5 Quality Control

Quality control of the neuroimaging data (*n*=203 scans) was completed, including manual inspection of each participant’s brain extraction, segmentation, registrations, masks, fibre directions, tracts, SC matrices, cleaned fMRI ROI time series and FC matrices (Figure 1).

**Figure 1.**
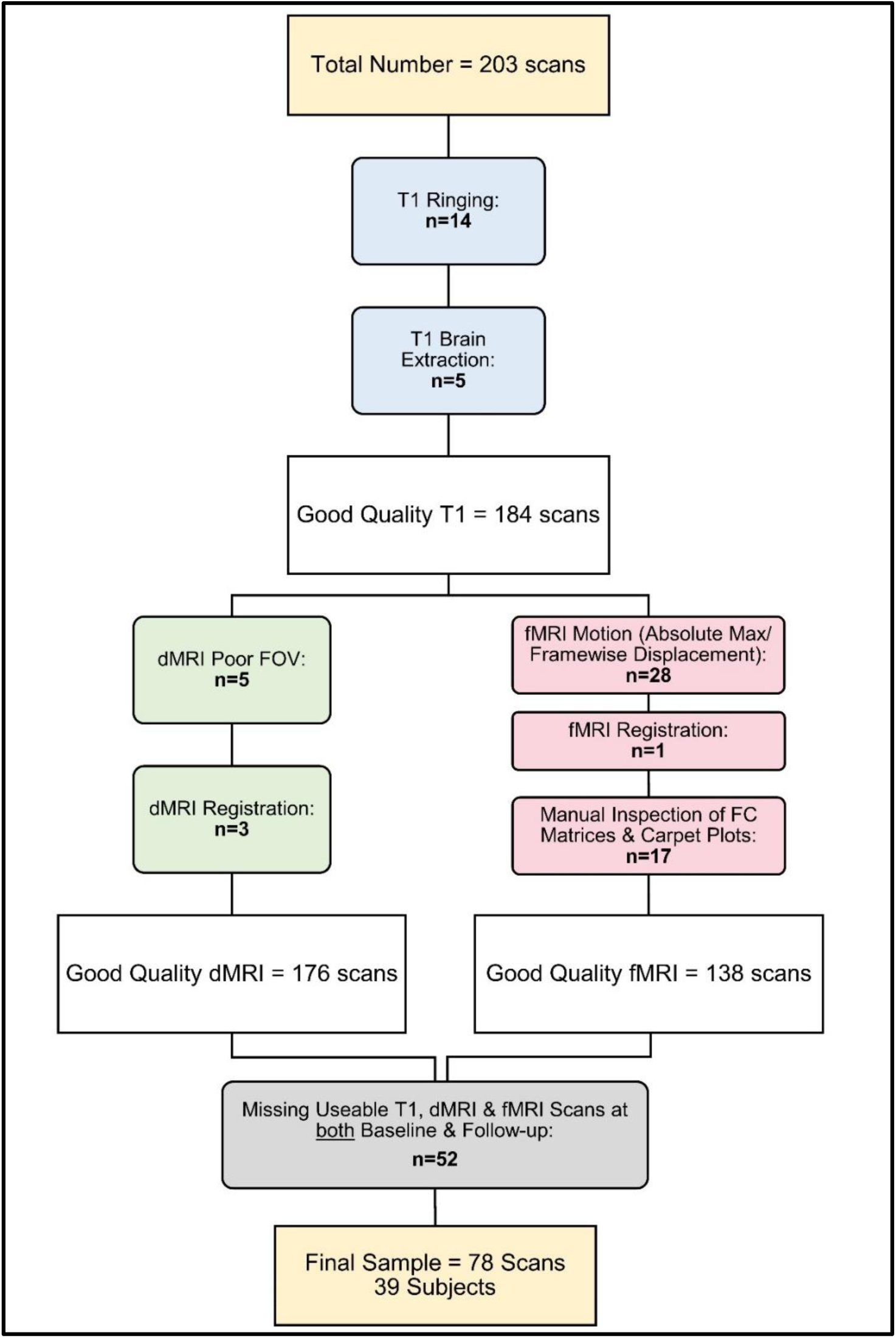
Exclusion and inclusion criteria. The initial dataset included 203 scans. The respective number of excluded scans are documented in bold. After quality control of the T1 images (blue), there remained 184 good-quality T1 images. The dMRI (green) and fMRI (red) scans were then assessed, resulting in 176 and 138 good-quality scans, respectively. Lastly, subjects without good quality T1, dMRI, and fMRI scans at both baseline and follow-up (i.e., three usable scans at each time point) were excluded from the final analyses. The resulting final sample included 39 subjects with scans at two-time points, totaling 78 scans. Abbreviations: FOV = field of view; FC = functional connectivity.

Participants’ T1 images with the presence of extreme motion artefacts (i.e, ringing) and poor brain extraction were noted and excluded (*n*=14 and *n*=5, respectively) with consideration of downstream functional and diffusion sub-pipeline outputs (e.g., the impact on registration). Additionally, scans with extremely poor FOV (e.g., missing additional regions to those excluded at the group level, see above) were excluded (*n*=5).

Scans with poor dMRI-to-T1 registration were also excluded (*n*=3). The SC matrices, along with the distribution of weights and tract lengths, were inspected for extreme sparsity. No other participants were excluded based on their SC matrices.

FMRI scans with greater than 8mm maximum absolute displacement or greater than 0.3mm framewise displacement for more than 8 minutes were excluded (*n*=28). Participants with extremely poor fMRI-to-T1 registration were noted and in conjunction with inspection of their FC matrices were excluded (*n*=1). More specifically, the FC matrices were checked for weak homotopic connectivity, indistinguishable intra-and inter-hemispheric quadrants and significant “banding” (i.e., an indication of motion artifacts or poor registration). The impact of residual motion artifacts was assessed with the MCFLIRT displacement plots and carpet plots of the ROI time series after cleaning. Scans with extreme residual motion based on the manual inspection of the FC matrices and carpet plots were excluded (*n*=17).

Lastly, participants who did not pass QC on all three neuroimaging data types (T1, fMRI, dMRI) for both a baseline and one-year follow-up visit were excluded (*n*=52), ensuring an equal number of subjects in each condition (i.e., a requirement for subsequent analyses described in Sections 2.7). Therefore, a total of 125 scans were excluded after all QC checks, resulting in a final analysed sample of 78 scans from 39 participants.

A number of additional QC assessments were completed on the final sample (*n*=78). Reports from the eddy current correction step (EDDY) indicated that the final analysed scans had no more than 3.5% total outliers (e.g., from motion causing signal dropout). After fMRI FIX cleaning, the temporal signal-to-noise ratio (tSNR) was not significantly different between time points, *t*(38)=-1.39, *p*=0.17. A linear mixed-effects model with participants’ post-FIX tSNR values and random intercepts for each participant (i.e., to account for within-participant variations) also found no significant relationship with either handedness, age, mean relative head motion or mean absolute head motion (*p’s*>0.26). For the final sample, there were also no significant differences in the mean absolute head motion between time points for the dMRI, *t*(38)=2.02, *p*=0.77, or fMRI data, *t*(38)=2.02, *p*=0.20. In addition, there were no significant differences in the mean relative head motion between time points for the dMRI, *t*(38)=-1.20, *p*=0.24, or fMRI data, *t*(38)=0.65, *p*=0.52. A QC-FC metric, the percentage of FC edges that correlated with head motion (average framewise displacement; *p*<0.05 uncorrected, Ciric et al., 2017; Graff et al., 2022a; Parkes et al., 2018), was calculated to be 16.8% (median *R*=0.075) for the final sample.

#### 2.4.6 Fingerprinting

As an indication of whether the adopted preprocessing cleaning approach of the functional data decreased or enhanced individual-specific information, we used a fingerprinting match rate approach described previously (Graff et al., 2022a). This metric was computed as the number of times an fMRI scan had the highest correlation with a second scan from the same participant, divided by the total number of scans (N=138). We calculated the fingerprinting match rate on all high-quality fMRI scans, including scans that did not have two high quality time points. These scans served as potential false matches. The match rates as percentages for the scans after preprocessing compared to no confound mitigation were 82.22% and 11.11%, respectively. Graff et al. (2022) achieved a maximal match rate of 90.2% on an overlapping sample.

### 2.5 Network Analysis

The Brain Connectivity Toolbox (BCT; https://sites.google.com/site/bctnet) was used to calculate the metrics for network analysis. Modularity, local clustering, and average weighted degree were chosen to characterise the network topology (e.g., segregation) in the developing brain.

The matrices’ diagonal (i.e., self-connections) were set to zero and negative weights were retained in the FC matrices. Modularity (Q) was calculated using the “community_louvain.m” function. Asymmetric treatment of negative weights was used for the FC matrices, such that a greater contribution was given by the positive weights (Rubinov and Sporns, 2011). Subjects’ Q values were also compared to 1000 null models that were generated with the “null_model_und_sign” function (i.e., preserved weight and degree distribution, approximated strength distributions) for each participant. For all participants, each individual’s SC and FC networks were significantly more modular than the null models (*p*’s<0.001), indicating that the modules identified are statistically different from chance and therefore meaningful to analyse. In order to assess changes in modularity with age, a linear mixed-effects model with participants’ SC modularity statistic was tested with age, handedness, sex, dMRI absolute and relative motion as fixed effects and random intercepts for each participant. Similarly, a model for participants’ FC modularity statistics with age, sex, fMRI absolute and relative mean displacement was also tested.

The local clustering coefficient is the ratio of the connections between a region’s neighbours (i.e. observed triangles) and the total possible number of such connections (Watts and Strogatz, 1998), and was calculated with the “clustering_coef_wu.m” (SC) and “custering_coef_wu_sign.m” (FC, Constantini & Perugini’s generalisation) functions. Subjects’ local clustering coefficient values were also compared to 1000 null models (generated with the “null_model_und_sign” function) for each participant. The local clustering was significantly greater than the null models (*p’*s<0.05) for 98.6% of the 15,054 (193 regions x 78 subjects) SC network regions and in 96.8% of the FC network regions.

For each participant’s SC matrix, average “weighted degree” was also calculated by taking the column-wise *average connection strength* for each region, resulting in a 1×193 SC vector (where 193 is the number of regions) (Rubinov and Sporns, 2010; Zimmermann et al., 2016). This was also completed for the participant’s FC matrices.

### 2.6 SC-FC Coupling

A SC-FC coupling metric was calculated as the Spearman’s correlation between a region’s SC and FC weighted degree vectors for each participant at each time point, following the approach described by Zimmerman and colleagues (2016). This resulted in a single SC-FC coupling value for each region of interest.

### 2.7 Partial Least Squares (PLS) Analyses

Partial Least Squares (PLS) correlation analyses (Krishnan et al., 2011; McIntosh and Lobaugh, 2004) were conducted to identify maximal covariance patterns in the data. PLS is particularly advantageous as it is able to cope with collinearity among the variables. Specifically, two condition mean-centred task PLS analyses and behavioural PLS (bPLS) analyses were conducted.

Mean-centred task PLS uses a mean-centred matrix calculated from the brain observations (specified as X) with two conditions (i.e., visits). This ‘task’ PLS identifies within-subject brain changes across the one-year follow-up, i.e. a longitudinal analysis.

Behavioural PLS analysis first computes a correlation matrix between a particular brain measure (specified as X) and the behavioural measures (specified as Y). Mutually orthogonal latent variables (LVs) are produced by decomposing the calculated data matrix using singular value decomposition (SVD). Each LV has a singular value and weights for the X and Y variables (i.e., brain and task/behaviour saliences, respectively). The bPLS identifies whether the brain-behaviour correlations are similar at time points one and two, i.e. a cross-sectional analysis that includes two time points assessed in parallel.

#### 2.7.1 Matrices for PLS analyses

The brain measures analysed were the modularity statistic, local clustering coefficients, regional average connection weights (i.e., weighted degree) and SC-FC coupling values. For a given region-wise brain measure, the vectors for each participant at time point one were stacked resulting in a matrix with dimensions 39 participants x 193 regions. This was also done for the vectors for participants at time point two. The resulting matrix for time point two was appended to the time point one matrix, resulting in a 78 scan (i.e., 39 participants x 2 scans) by 193 region matrix. The same steps were completed for each network metric and the SC-FC coupling values. Mean-centred task PLS analyses were conducted for each brain measure between the two visits. Behavioural PLS analyses were performed for each brain measure with age and attention measures.

An additional bPLS analysis was conducted with the SC and FC weighted degree, as well as the SC-FC coupling (at baseline and follow-up). The SC, FC, and SC-FC matrices were all of dimensions 78 scans x 193 regions and were stacked to create the resulting brain data matrix (234 scans x 193 regions).

#### 2.7.2 Tests of Specificity

The cosine similarity of the brain saliences from each respective task PLS analysis and bPLS were calculated to identify whether the regions that were different between time points (i.e., identified by the task PLS) were the same as those that were correlated with age and behaviour (i.e., identified by the bPLS). This was meant as a qualitative assessment only.

Furthermore, separate two-condition bPLS analyses were conducted with each brain metric with sex and motion metrics (i.e., mean relative fMRI motion and percentage of dMRI outliers for FC and SC metrics, respectively). Both motion metrics were used for the SC-FC coupling metric analyses. The cosine similarity of the brain saliences from each PLS analysis and the respective bPLS analyses with sex and motion were calculated. This was conducted to assess whether the regions identified by the task PLS and bPLS analyses with attention and age were different from those that were correlated with sex and motion. Permutation testing (i.e. resampling without replacement) was completed for each respective analysis (1000 permutations) to test the significance of the cosine similarity values.

#### 2.7.3 Significance, reliability and reproducibility tests

A total of 1000 permutations were conducted to test the significance of the PLS results. As a reliability test of the weights within each LV, 500 iterations of bootstrapping were conducted. The number of permutations and bootstrapping were determined according to the standards in the literature (Efron and Tibshirani, 1994; Marozzi, 2007). Importantly, bootstrapping suggests that the features of the connectomes assessed are robust. Bootstrap ratios (BSRs) are used to assess the contribution of particular regions or edges to the overall effect. They are interpreted as a reliability score, similar to z-scores where a value of 2.0 corresponds approximately to a 95% confidence interval. BSRs greater than 2.0 therefore indicate the brain regions that reliably contribute to the latent variable.

Recently, it was demonstrated that performing null hypothesis testing for statistical significance does not assure reproducibility of the results (unpublished observations). Therefore, two additional assessments of the reproducibility of the singular values and singular vectors across various sample sizes were conducted (“test-train” and “split-half”, respectively). A “test-train” assessment was performed such that the analysis was performed on random halves of the original sample (split-half resampling) with subjects’ time points held together. The assessment was completed to test if the same LV pattern can produce a similarly strong covariance between X and Y matrices (i.e., indicated by the singular value).

Thus, 500 resamples were performed to produce a distribution of test *<u>singular values</u>*. The distribution’s mean and standard deviation were used to compute a z-score whereby a larger value indicates greater reproducibility of the singular values. Secondly, the “split-half” LV reproducibility test was performed in the same way as above (500 random split-half resamples) but the *<u>similarity of singular vectors</u>* was computed. For both assessments, the calculated z-scores (from the singular value and singular vector distributions, respectively) are also compared to the corresponding z-scores computed from null distributions generated by permutation resampling (i.e., difference greater than ±2).

#### 2.7.4 Code Accessibility

The code (run on macOS Big Sur Version 11.5.2) for the computation of the assessed metrics, PLS analyses, and visualisation of the key figures are freely available online at https://github.com/McIntosh-Lab/Rokos2024_SCFC_NetworkAnalyses.

## 3. Results

### 3.1 Attention Measures

The paired-sample t-tests indicated significant performance improvements for each attention task between time points (Table 1). The mixed-effects further indicated that age was significantly positively associated with sustained attention (*β*=2.30, *SE*=0.33, *t*=6.99, *p*<0.0001), selective attention, (*β*=1.78, *SE*=0.29, *t*=6.14, *p*<0.0001), and executive attention (*β*=4.03, *SE*=01.23, *t*=3.28, *p*=0.0016). The mixed model results are also reported in Extended Data Table 1-1.

**Table 1.** T-tests for attention measures between initial and follow-up visits.

| Measure | Time point 1<br>Mean (SD) | Time point 2<br>Mean (SD) | t-test(df) | p-value |
| --- | --- | --- | --- | --- |
| Sustained<br>Attention | 23.94(3.79) | 26.78(2.82) | 6.48(38) | $p<0.0001$ |
| Selective<br>Attention | 14.15(3.16) | 15.9(2.45) | 3.64(38) | $p<0.001$ |
| Executive<br>Attention | -33.79(13.22) | -27.62(6.58) | 2.80(38) | $p=0.0079$ |

### 3.2 Longitudinal Change Assessments

#### 3.2.1 Mean-Centred Task PLS

In order to explore within-subject changes over the follow-up period, mean-centred task PLS analyses were conducted for each brain measure. The modularity statistics did not change significantly between the baseline and follow-up visits for the SC (*p=*0.09) or FC matrices (*p=*0.05). Additionally, there were no significant longitudinal changes in any of the region-wise SC local clustering (*p=*0.23) or SC weighted degree (*p=*0.12) metrics.

The region-wise FC graph theory metrics exhibited significant alterations over the year, including reduced FC local clustering (*p*=0.048, *Z*_test-train_=1.89,*Z*_null_test-train_=0.096; *Z*_split-half_ =2.71,*Z*_null_split-half_=1.45) and weighted degree (*p*=0.026, *Z*_test-train_=2.50; *Z*_null_test-train_=0.025; *Z*_split-half_ =2.94; *Z*_null_split-half_=1.39). Several regions showed a decrease in FC local clustering and weighted degree, while the right medial PFC exhibited increases in both metrics (Figure 2 A&B). Based on the reproducibility tests, only the weighted degree result was reproducible based on the test-train assessment (i.e., Z value exceeding the Z_null_ value by more than two).

**Figure 2.**
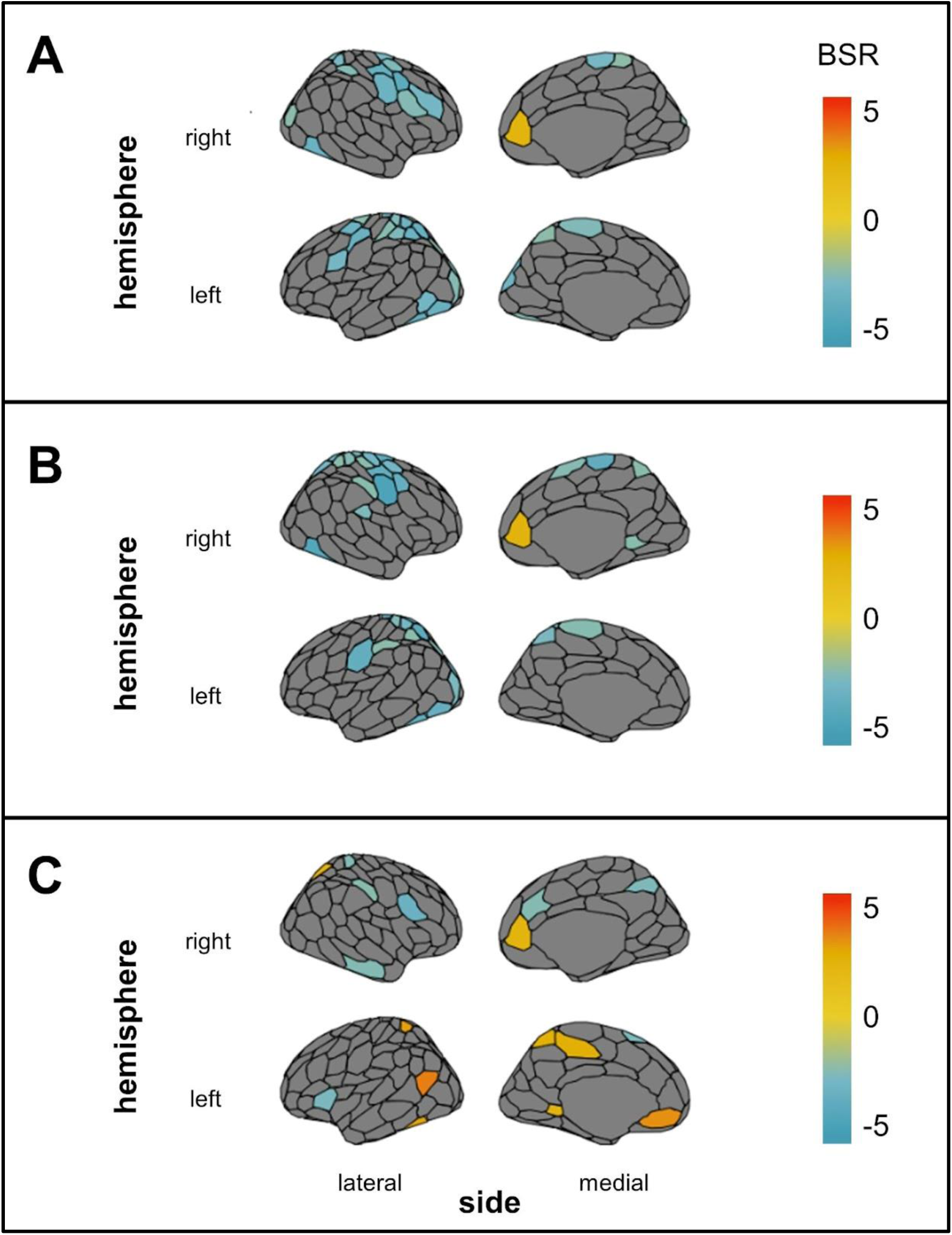
Mean-centred task PLS results. Over the one-year follow-up period, bootstraps ratios (BSRs) indicate regionally specific decreases (negative BSRs, green/blue) and increases (positive BSRs, yellow/red) in functional connectivity (FC) local clustering (***A***), FC weighted degree** (***B***), and the SC-FC coupling metric (***C***). BSRs thresholded at +2.0 and -2.0 are plotted onto a brain. Analyses that passed the reproducibility tests are marked with **. Abbreviations: BSR = bootstrap ratio; structural connectivity = SC; functional connectivity = FC.

The analysis with the SC-FC coupling metric identified one significant LV (*p*=0.014, *Z*_test-train_=-0.19;*Z*_null_test-train_=-0,17; *Z*_split-half_ =1.32;*Z*_null_split-half_=1.30), indicating a regional pattern of increasing (e.g., left and right medial PFC, right superior parietal lobule, left postcentral gyrus), and decreasing (e.g., right lateral PFC, left insula) coupling between SC and FC (Figure 2C). The reproducibility tests indicated that this relationship was not reproducible.

Extended Data Table 2-1 documents the ROI name abbreviations for the ROIs assessed for all region-wise analyses. The cosine similarities between the brain saliences for each task PLS analysis with the respective bPLS analyses with sex and motion, as well as the associated *p*-values are reported in Extended Data Table 2-2. The cosine similarities and permutation testing indicated that there was little overlap between the regions identified by the mean-centred PLS analyses and those associated with sex or motion.

#### 3.2.2 Age & Longitudinal Changes

For the community statistics (Q), linear mixed-effects models were also conducted to identify whether changes in modularity were associated with age (with participants as the random intercept). The subject’s handedness, sex, and the dMRI head motion metrics were included to ensure the effects of motion were controlled for. The model indicated that age was negatively associated with SC modularity (*β*=-0.40, *SE*=0.11, *t*=-3.79, *p*=0.0003, Figure 3A). The estimated variance by the participant random intercepts was -0.13(*SD*=0.59). FC modularity was not significantly associated with age (*p*=0.15).

**Figure 3.**
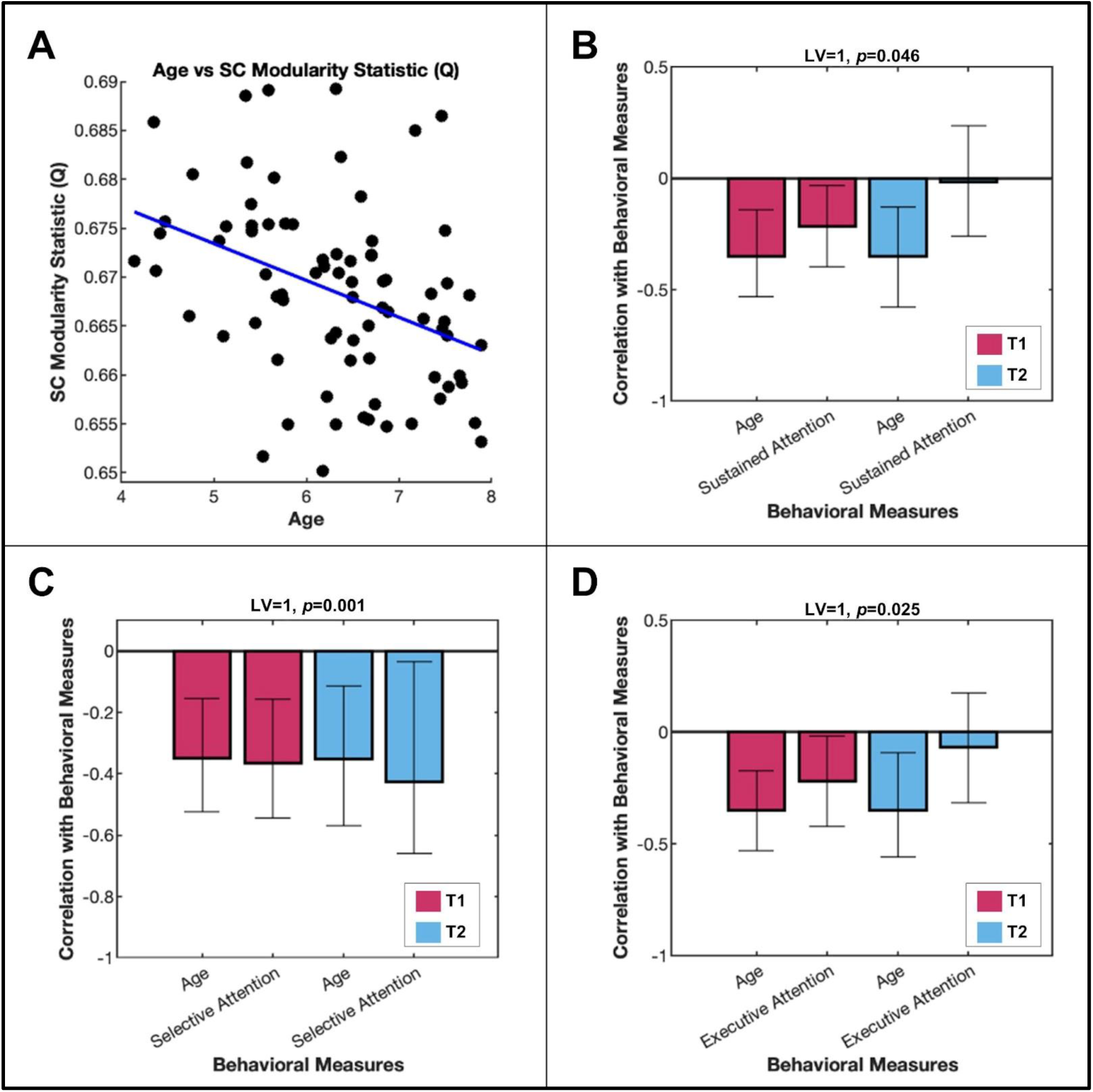
Structural connectivity modularity results. Structural connectivity (SC) modularity was negatively associated with age (***A***), as well as sustained attention** (***B***), selective attention** (***C***), and executive attention** (***D***) across both the initial and follow-up visits. Analyses that passed the reproducibility tests are marked with **. Abbreviations: LV = latent variable; T1 = time point one; T2 = time point two; structural connectivity = SC.

For each significant mean-centred PLS analysis, the correlation between the differences in participants’ brain scores (between time points) with their baseline age were calculated. The correlations were conducted to assess whether the significant longitudinal changes in region-wise metrics (i.e., the region-wise FC metrics, SC-FC coupling) were associated with baseline age. There were no significant correlations between the differences in brain scores and age (|*r|*’s<0.18, *p*’s>0.28).

### 3.3 Cross-Sectional Assessment - Behavioural PLS

#### 3.3.1 SC Modularity

The bPLS analysis between the SC modularity statistic (Q) with age and sustained attention revealed a significant LV (*p*=0.046; *Z*_test-train_=2.35;*Z*_null_test-train_=-0.08; *Z*_split-half_ =4.39;*Z*_null_split-half_=2.21), whereby lower modularity was associated with older age and better sustained attention at time point one. The SC modularity statistic was also negatively associated with age and selective attention at both timepoints (*p*=0.001; *Z*_test-train_=3.17;*Z*_null_test-train_=0.08; *Z*_split-half_ =8.59;*Z*_null_split-half_=2.61), and executive attention at time point one (*p*=0.025; *Z*_test-train_=2.72;*Z*_null_test-train_=-0.041; *Z*_split-half_ =4.31;*Z*_null_split-half_=2.14).

The test-train and split-half resampling tests indicated that these results were reproducible, with Z values exceeding the Z_null_ values by more than two. The modularity PLS analyses are visualised in Figure 3B-D.

#### 3.3.2 SC Local Clustering

Behavioural PLS analyses for SC local clustering with age and sustained attention identified one significant LV (*p*<0.001; *Z*_test-train_=0.21;*Z*_null_test-train_=-0.02; *Z*_split-half_ =1.55; *Z*_null_split-half_=1.39). For older participants, worse sustained attention was associated with lower structural local clustering for regions in the default mode network (e.g.,left and right dorsal PFC, right ventral PFC) (Figure 4A, negative BSRs), as well as greater local clustering for the right superior parietal lobule (SPL), left somatomotor cortex, and extrastriate regions.

**Figure 4.**
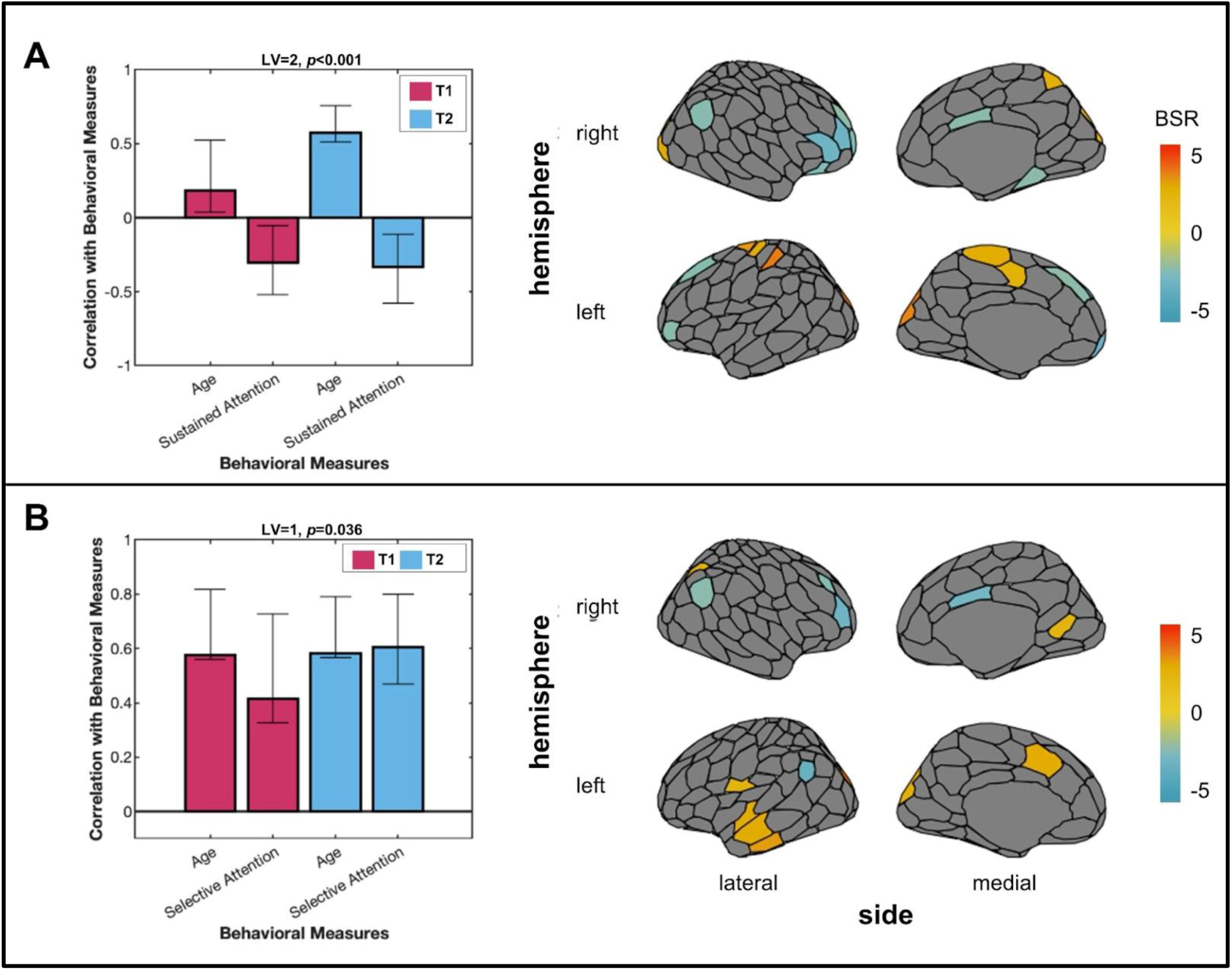
Structural connectivity local clustering behavioural PLS results. ***A***, The significant latent variable for the behavioural PLS between structural connectivity (SC) local clustering with age and sustained attention revealed an age-dependent association. ***B***, The significant latent variable for the analysis between SC local clustering with age and selective attention revealed that SC local clustering was positively associated with age and selective attention at both time points. Bootstrap ratios thresholded at +2.0 and -2.0 are plotted onto a brain. Abbreviations: BSR = bootstrap ratio; LV = latent variable; T1 = time point one; T2 = time point two; structural connectivity = SC.

Furthermore, one significant LV was identified for SC local clustering, age, and selective attention (*p*=0.036; *Z*_test-train_=0.17;*Z*_null_test-train_=-0.07; *Z*_split-half_ =1.44;*Z*_null_split-half_=1.31) (Figure 4B). Regions with greater local clustering, such as the left and right inferior parietal lobule and dorsal PFC were negatively correlated with selective attention and age, at both time points (negative BSRs). Local SC clustering for regions such as the left and right extrastriate cortex, right SPL, and left temporal cortex were positively correlated with age and attention (positive BSRs).

The analysis with SC local clustering, age, and executive attention did not identify any significant LVs (*p*=0.11). The cosine similarities between the brain saliences from each SC local clustering bPLS (with age and attention), and the respective task PLS analysis and bPLS analysis with potential confounds were computed. The permutation testing for each respective analysis indicated that there was little overlap between the regions that were correlated with age and behaviour and those associated with sex or motion (Extended Data Table 4-1).

#### 3.3.3 SC Weighted Degree

The SC weighted degree PLS with age and sustained attention revealed two significant LVs. The first LV (*p*=0.013; *Z*_test-train_=0.013;*Z*_null_test-train_=0.0098; *Z*_split-half_ =1.32; *Z*_null_split-half_=1.40) identified regions where weighted degree was positively correlated (e.g., left precuneus, right intraparietal sulcus, right superior and inferior parietal lobule; indicated by positive BSRs), as well as regions where weighted degree was negatively correlated (e.g., left somatomotor cortex, right lateral and dorsolateral PFC; indicated by negative BSRs) with age and sustained attention (Figure 5A). The second LV (*p*=0.001; *Z*_test-train_=0.25;*Z*_null_test-train_=-0.06; *Z*_split-half_ =1.24; *Z*_null_split-half_=1.32), revealed a unique set of regions where poorer sustained attention was correlated with SC weighted degree for older children. For instance, worse sustained attention in older children was associated with greater weighted degree (positive BSRs) for regions associated with visual processing (e.g., left and right extrastriate), and lower weighted degree (negative BSRs) for regions in the default mode network (e.g., left and right medial PFC, left dorsal PFC) (Figure 5B).

**Figure 5.**
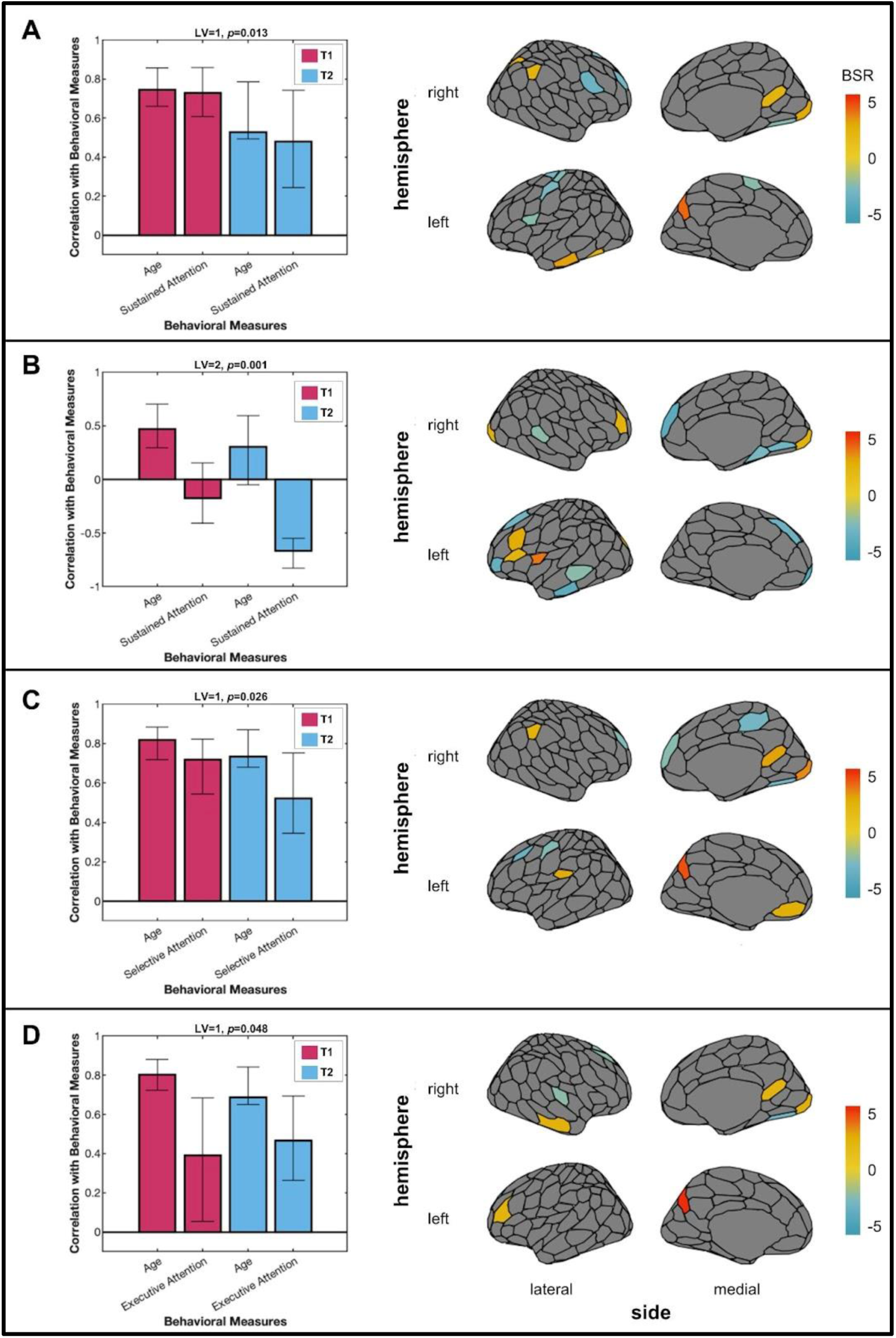
Structural connectivity weighted degree behavioural PLS results. ***A***, The positive association between structural connectivity (SC) weighted degree with age and sustained attention, for both time points was evidenced by the first latent variable. ***B***, A different pattern of brain regions where the relationship between SC weighted degree and sustained attention depended on age was revealed by the second latent variable. The first latent variables for the behavioural PLS analyses between the SC weighted degree with age and selective attention (***C***), and executive attention (***D***) also showed a positive association between SC, age, and attention. Bootstrap ratios thresholded at +2.0 and -2.0 are plotted onto a brain. Abbreviations: BSR = bootstrap ratio; LV = latent variable; T1 = time point one; T2 = time point two; structural connectivity = SC.

The significant LV which indicated a significant association between SC weighted degree with age and selective attention also revealed a distinct pattern of brain regions at both time points (*p*=0.026; *Z*_test-train_=-0.03;*Z*_null_test-train_=-0.02; *Z*_split-half_ =1.32;*Z*_null_split-half_=1.28) (Figure 5C). Specifically, greater weighted degree was associated with better selective attention in older children in regions such as the left precuneus, left secondary somatosensory cortex, and right superior and inferior parietal lobule. Weighted degrees for a number of prefrontal regions (e.g., left and right lateral PFC, right medial PFC and dorsal PFC) and medial parietal cortex were negatively correlated with age and selective attention.

An LV indicated a significant association between SC weighted degree with age and executive attention (Figure 5D) independent of the time point (*p*=0.048; *Z*_test-train_=-0.50;*Z*_null_test-train_=-0.004; *Z*_split-half_ =1.24; *Z*_null_split-half_=1.30). In this LV, the left precuneus, left lateral ventral PFC, right striate, right temporal cortex, right retrosplenial cortex, and right SPL were positively correlated with executive function and age, while the right dorsal and dorsolateral PFC, extrastriatal cortex, auditory cortex, and left dorsal PFC were negatively correlated.

The cosine similarities between the brain saliences for the SC weighted degree bPLS analyses (with age and attention), with the respective task PLS analysis and bPLS analysis with sex and motion, indicated that there was also little overlap between the identified regions (Extended Data Table 5-1).

#### 3.3.4 FC Graph Theory Metrics

A single significant LV for the FC graph theory metric weighted degree was identified with age and selective attention (*p*=0.038), however, the relationship was unstable and weak (i.e., confidence intervals crossed zero, correlations<0.22). No other significant LVs were observed between the FC metrics, age, and any of the attention measures (*p*’s>0.08). The cosine similarities between the brain saliences for each FC graph theory bPLS analyses and the respective task PLS analyses indicated that there was little overlap in the regions that differed between the two time points and the regions correlated with age and attention (*p*’s>0.12). Similarly, the cosine similarities between the brain saliences for the FC bPLS analyses (with age and attention) with the respective bPLS analyses with sex and motion also indicated that there was little overlap between the regions that were correlated with age and behaviour and those associated with sex or motion (*p*’s>0.15).

### 3.4 SC-FC Coupling With Age & Attention Measures

Next, we wanted to explore whether the coupling between SC and FC would collectively identify important brain regions across development. A bPLS analysis was run with SC and FC weighted degree and SC-FC coupling as independent variables, and age as the dependent variable. The one significant LV (*p*=0.011, *Z*_test-train_=3.79;*Z*_null_test-train_=-0.05; *Z*_split-half_ =9.56; *Z*_null_split-half_=1.54) indicated that SC weighted degree was the dominant predictor of age (Figure 6). Additional bPLS analyses of only SC-FC coupling with age and each attention measure revealed no significant LVs (*p*’s>0.06).

**Figure 6.**
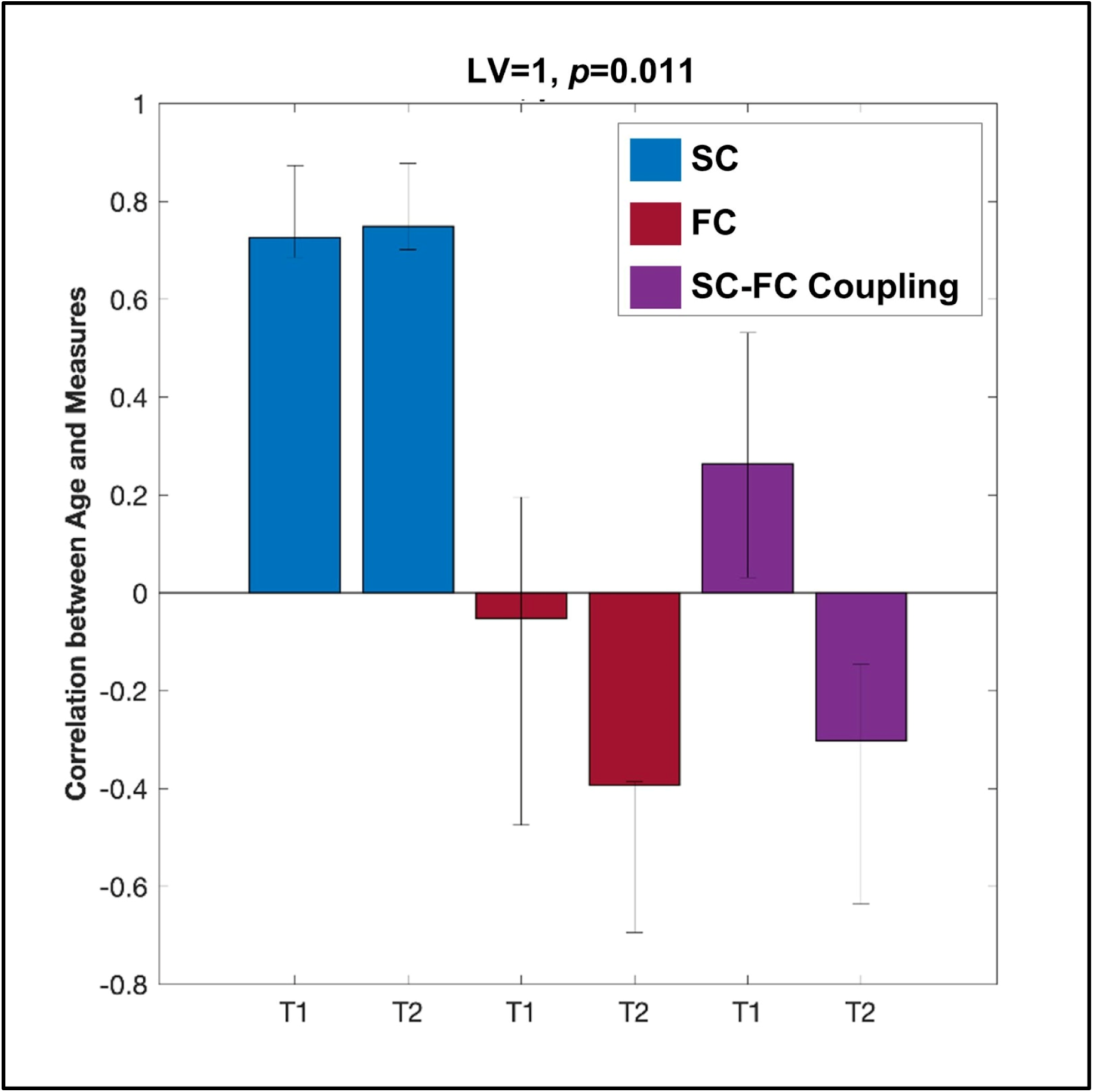
Combined behavioural PLS result with age. The correlation between structural connectivity (SC) weighted degree, functional connectivity (FC) weighted degree, and SC-FC coupling with age. The latent variable with confidence intervals on each of these correlations show that SC reliably and strongly predicts age, independent of time point. Abbreviations: BSR = bootstrap ratio; LV = latent variable; T1 = time point one; T2 = time point two; structural connectivity = SC; functional connectivity = FC.

The cosine similarities between the brain saliences for the SC-FC coupling PLS analyses with the respective task PLS analyses and bPLS analyses with sex and motion are reported in Extended Data Table 6-1. The regions identified by the significant bPLS analysis with SC weight degree, FC weighted degree, SC-FC coupling, and age did not overlap highly with those associated with sex or motion (*p*=0.35).

## 4. Discussion

This study utilised graph theory analyses to investigate the relationship between both structural connectivity (SC) and functional connectivity (FC) with age and attentional abilities during early childhood. Mean-centred task PLS analyses were used to assess longitudinal brain changes in typically developing children between the ages of 4 and 8, a relatively underexplored age-range. Behavioural PLS analyses were used to explore age-associations as well as whether brain network features were related to sustained, selective, and executive attention at each visit.

We identified an association between age and longitudinal developmental decreases in SC modularity (i.e., linear mixed-effects model), as well as associations of lower SC modularity with better attention performance (i.e., bPLS). Network metrics (i.e., local clustering, weighted degree) of the structural networks did not change significantly over the follow-up period but were associated with age and attention at both time points. The bPLS analyses found two distinct brain-behaviour patterns (i.e., latent variables). The first pattern found, across all three attention measures, regions where greater SC weighted degree was associated with older age and better attention, as well as where weaker SC weighted degree was associated with younger age and worse attention. Similarly, greater SC local clustering was associated with better selective attention for older children. The second pattern identified age-dependent relationships for SC local clustering and for SC weighted degree with *sustained* attention, where the regional SC metrics were associated with worse attention for older children but better attention for younger children.

On the other hand, regional FC graph theoretical metrics identified key brain regions that changed between baseline and the one year follow-up period but were not associated with age or attention. An additional analysis with SC and FC, as well as SC-FC coupling further revealed structural connectivity as a unique predictor of age during this developmental period. The evident relation between brain network features with attention highlights the opportunities for behavioural interventions or therapies between infancy and middle-to-late childhood (as discussed by Slattery et al., 2022).

### 4.1 Structural Connectivity

#### 4.1.1 SC Longitudinal Changes

The longitudinal analyses offer a view of brain ‘development’ specifically, beyond the associations with age that can be identified in cross-sectional studies but are at risk of bias (Di Biase et al., 2023). To improve our understanding of brain network development across early childhood, we explored network changes between baseline and the one-year follow-up period. In line with previous research, we also documented an age-related decrease in SC modularity, signifying a decline in global segregation or greater intermodule integration of structural networks (Cao et al., 2017; Dennis et al., 2013; Hagmann et al., 2010; Tymofiyeva et al., 2013). There were no significant longitudinal changes in the SC local clustering or weighted degree metrics, suggesting a more stable regional SC topology over the one year period investigated in this early childhood timeframe. The longitudinal results are further complemented by the bPLS results which characterised stable structural brain network-behaviour associations in early childhood.

#### 4.1.2 SC Cross-sectional Age & Attention Associations (bPLS)

Although a number of developmental studies have explored age-related differences in structural connectomes, few have assessed their relationship with behaviour during early childhood. In order to improve our understanding of how brain network topology is related to behaviour in this age range, we investigated the relationship of modularity with attention measures. We demonstrated that lower modularity was associated with better attention in older children. This complements clinical studies where children with ADHD (i.e., worse attention) have been found to exhibit greater modularity than controls and fewer between sub-network connections (Beare et al., 2017). Importantly, we also expanded upon the characterised global developmental topology of segregation that supports attention by investigating *nodal* metrics.

The latent variables for the regional network analyses identified key regions that were associated with age and attention at baseline and follow-up. The cross-sectional SC-behaviour relationship and age-associations were generally consistent for both time points, and is consistent with the longitudinal analyses that found no significant changes in SC regional metrics over the one-year. The results may be due, in part, to the short follow-up period and can be interpreted as a relationship that is stable or replicated at both visits. For instance, the SC local clustering analyses with selective attention and age, as well as the weighted degree analyses with all attention measures showed positive relationships at both time points. Nonetheless, the cross-sectional analyses allowed for the investigation of age- and attention-related associations over a longer time window (four years) as opposed to the one year follow-up period.

More specifically, reduced SC local clustering of the inferior parietal lobule (IPL) and prefrontal regions and greater local clustering of the right superior parietal lobule (SPL) was associated with older age and better selective attention. The temporal-parietal junction (including the IPL) has shown activation during stimulus-driven attention tasks, involving maintenance, orientation, and reorientation of attention (Igelström and Graziano, 2017). The observed pattern suggests that for older children, lower structural segregation of these regions supports better attention, but that segregation of the SPL may be favourable. The right SPL has previously been identified as a critical mediation region for visuospatial attentional tasks (Wu et al., 2016), therefore interventions that target the right SPL in early childhood may be particularly beneficial for selective attention performance in younger children.

The weighted degree measure also identified the right SPL as a key region where greater average SC strength was associated with better attention and older age. Although there was overlap in the identified regions between attention tasks (e.g., left precuneus, right SPL), SC weighted degree in unique medial and frontal regions were associated with age and selective attention (e.g., negatively associated with the right medial PFC and medial parietal cortex). Investigating the three different attention measures separately indicated that despite similarities in the regions associated with sustained, selective, and executive attention, they also had unique patterns. Additionally, only sustained attention had an age-dependent relationship with SC local clustering and weighted degree.

#### 4.1.3 SC Age-Dependent Relationships with Sustained Attention

The bPLS analyses identified unique patterns of brain regions where SC local clustering and weighted degree were associated with *worse* sustained attention for older children but *better* sustained attention for younger children. For example, worse sustained attention for older participants was linked to greater local clustering for a number of extrastriate cortex regions and left somatomotor cortex regions.

The SC weighted degree analysis also highlighted a distinct pattern of SC weighted degree associated with poorer sustained attention for older children, including greater weighted degree in visual processing regions and lower weighted degree for regions associated with the default mode network (DMN). These results align with early development of sensory regions (Reynolds et al., 2019; Stiles and Jernigan, 2010; Whitaker et al., 2016) and emphasise the significance of age-dependent, region-specific structural segregation for sustained attention performance. Our findings highlight an importance of visual sensory regions for younger children’s attention, while also indicating that the brain network’s continuing reliance on those regions may be detrimental for older children.

Furthermore, worse sustained attention for older children was associated with greater local clustering for dorsal attention network (DAN)-associated regions (e.g., right SPL and post-central gyrus), and lower local clustering for prefrontal DMN regions. These findings reinforce the significance of a network that incorporates diverse changes in both DAN and DMN regions (Esterman et al., 2013). Given the established associations of the DMN with mind-wandering and internally-directed processing, our findings suggest that less segregation of communication between the DMN and particular sensory and attention network regions for older children may negatively impact attention. The results indicate that the potential beneficial or deleterious influence of specific regions in structural networks is dependent on age, and only during a specific window of development, once again underscoring the need for individualised interventions. Moreover, understanding these intricate network changes may yield valuable insights into neurodevelopmental conditions like ADHD, where the balance between small-world properties of DMN, DAN, and visual network regions appear to be disrupted (Soman et al., 2023).

In summary, two latent variables were identified between SC weighted degree, age, and sustained attention. The latent variables characterised an intricate association for older children, where worse sustained attention was linked to greater SC weighted degree in primary visual regions, while better sustained attention was associated with greater SC weighted degree in integration regions like the SPL. The findings emphasise the relevance and relationship between behaviour and the reorganisation of structural networks that supports specific regions and networks (e.g., SPL, DAN, DMN) across development. This research underscores the significance of structural network changes in shaping childhood attentional abilities and their potential relevance for addressing age-specific, attention-related challenges in clinical contexts.

### 4.2 Functional Connectivity

#### 4.2.1 FC Longitudinal Changes

In addition to SC graph theory metrics, we explored how children’s functional networks changed over the one-year follow-up period. We found that the FC local clustering and weighted degree decreased over the year in numerous visual processing regions and regions associated with the DAN. Interestingly, only the medial PFC exhibited increased local clustering and weighted degree, suggesting its increasing role as a network hub (Hodel, 2018). Cross-sectionally, age-related FC decreases between the DAN and DMN in typically developing children have been reported (Rohr et al., 2017). An increase in self-stability of the functional connectomes over the one-year follow-up has also been reported to drive greater connectome individualisation with age (Graff et al., 2022b). Therefore, along with changes across the follow-up period, we investigated how age was related to FC network development.

On a regional level, the longitudinal changes in the FC metrics were not associated with children’s baseline age, indicating a consistent developmental pattern of change across this narrow age-range. In line with this, for the bPLS analyses (discussed further below), the FC metrics had no stable cross-sectional brain-behaviour relationships and age-associations. The identification of only longitudinal changes for FC regional metrics may be attributable to FC demonstrating larger variability between subjects. To explore this, we examined how the variability between the structural and functional connectivity matrices differed across subjects via principal component analyses (PCA). The PCA analyses identified that both SC and FC had relatively large first components, indicating a substantial proportion of variance that was common across subjects. Additionally, FC exhibited a smaller first component and subsequent components explained a larger portion of variance in the FC compared to SC (Extended Data Table 6-2). These differences in variance between SC and FC have previously been observed in an ageing population (Zimmermann et al., 2019), and highlights the advantages of longitudinal analyses that may be more sensitive to within-subject changes over time. Overall, we characterised a developmental pattern of functional brain network changes over the one-year follow-up period with no observed functional network-behaviour associations in early childhood.

#### 4.2.2 FC Cross-sectional Age & Attention Associations (bPLS)

The FC graph theory metric analyses found no significant or reliable attention- or age-associations. Further investigation with a wider age range would support the characterization of potentially more prominent age-related differences. Rohr and others (2018) identified distributed functional network integration across this age range, but compared to the current study focused on graph theory metrics, their study used analyses of specific networks (i.e., independent component analysis) in a larger sample (Rohr et al., 2018). Future work should also include analyses that capture more specific within-subject changes in brain-behaviour relationships (e.g., mixed-effects models), as well as the non-linear developmental trajectories that have been characterised in a number of functional network studies (Lebel et al., 2012, 2008). Additionally, dynamic FC has been proposed to be more sensitive to unique aspects of behaviour compared to static measures of FC (Eichenbaum et al., 2021). Therefore, characterising how both static and dynamic FC changes relate to attentional performance across early childhood would allow for an improved understanding of the neural dynamics in the developing brain (Naik et al., 2017). Nonetheless, our findings emphasise the importance of the structural connectivity between the functional networks associated with attention processing during childhood. The results of this study suggest SC topology may play a more substantial, underlying role in attention performance, considering the outsized effect of SC compared to FC and SC-FC coupling in the combined analysis.

### 4.3 SC-FC Coupling

The SC-FC coupling metric was found to change over a year of development, with a region-specific pattern of increasing and decreasing coupling between SC and FC. While the findings highlight the dynamic nature of brain development in early childhood, the changes in coupling were found to have low reproducibility and were not associated with participants’ age. The differences in between-subject variance between structural and functional connectivity has also been suggested to limit the subject specificity of the correspondence between SC and FC (Zimmermann et al., 2019).

Similarly, the cross-sectional bPLS analyses between the SC-FC coupling metric with age and each attention measure were non-significant. This lack of association might also be attributable to the narrow age-range of the current sample. Prior research found that the relationship between SC and FC stabilises and strengthens with age, however these studies considered older and larger age ranges (Betzel et al., 2014; ages 7-85), as well as fewer subjects (Hagmann et al., 2010; *n*=14, ages 2-18). Furthermore, the current SC-FC coupling approach considered all connections, including those supported by indirect SC (Honey et al., 2009). This is a distinction from literature that has focused solely on connections with direct SC links (e.g., Soman et al., 2023).

This study further investigated SC-FC coupling in tandem with SC and FC (weighted degree) and identified structural connectivity as the critical predictor of age. Although the SC-FC coupling metric used has previously been shown to reliably predict age for individuals between the ages 18 to 82 (Zimmermann et al., 2016), the younger sample assessed here may suggest a more distinct independence between SC and FC in early childhood. Another important consideration is that changes in SC appear to precede FC changes (Wendelken et al., 2017; Zuo et al., 2017). This may suggest the possibility of a lead-lag association where SC at early time points predicts changes in later FC and behaviour (Wendelken et al., 2017). Although changes in SC likely play a role in constraining and shaping functional networks, spontaneous and experience-dependent neuronal activity is also thought to influence the development of brain networks (Deco et al., 2013; Fair et al., 2007; Honey et al., 2009; Mišić et al., 2016; Wang et al., 2013). Therefore, computational modelling is a valuable approach that may allow for a better understanding of how SC and FC are causally related to each other in this developmental period.

Computational frameworks (e.g., The Virtual Brain; Sanz Leon et al., 2013) that integrates multiple neuroimaging modalities (e.g., both SC and FC) and non-linear network dynamics (e.g., dynamic FC) for typically developing children would provide more extensive insights into healthy brain network trajectories of maturation.

### 4.4 Important Considerations & Limitations

The test-train and split-half resampling approaches described in this study emphasise the importance of considering the level of reproducibility, in addition to the reliability (bootstrapping) and significance (permutation testing) of each analysis (Nakua et al., 2023). For example, a number of significant brain-behaviour relationships were not reproducible, likely due to the small sample size analysed. In order to improve the generalizability of the findings, future work should consider data collected from multiple sites, a larger sample, wider age range, and follow-ups beyond one year. The smaller sample size analysed was in part due to rigorous data processing/cleaning and quality control to mitigate the potential impact of motion (Grayson and Fair, 2017; Oldham and Fornito, 2019; Power et al., 2012).

The passive-viewing functional scans may be confounded by the level of attentiveness and enjoyment of the show during the scan, which would add more intersubject variability (e.g., Rohr et al., 2016), therefore subsequent studies should consider including a measure to assess participants’ level of engagement. As discussed previously (Rohr et al., 2018), the behavioural attention measures may also be influenced by situational context (e.g., children may prioritise their attention differently depending on the context) and may be limited in terms of real-world applicability. Lastly, the current parcellation only includes cortical regions (due to the likelihood of increased false positive connections within subcortical regions), however atypical fronto-subcortical connectivity has been displayed in children with ADHD (Kumar et al., 2022; Rosch et al., 2023). In order to achieve a more comprehensive understanding of the interplay between the whole-brain structure-function relationship with attention, future work should explore whether the results are impacted by the inclusion of subcortical regions and the use of different parcellation approaches.

### 4.5 Conclusion

In children aged 4 to 8 years, region-wise graph theory analyses supported the characterization of variable developmental changes and brain-behaviour relationships across the brain. Over a one-year follow-up period, longitudinal changes in functional graph theory metrics were observed, as well as age-related decreases in SC modularity. This study further emphasises how structural topology is related to age and attentional performance. The local clustering and weighted degree metrics identified key regions where lower SC segregation was associated with better selective attention skills in older children, but also differentiated regions (e.g., right SPL) where greater SC weighted degree and clustering appeared to be beneficial. Evidently, early childhood is an extremely dynamic period where cognitive functioning is intricately and predominantly linked to structural network features. The current findings carry numerous implications for understanding healthy development and identifying potential targets for neurodevelopmental disorders.

## Acknowledgements

This work was funded by the CIHR Project Grant, NSERC Discovery Grant, and CIHR-INMHA Bridge Grant. This research was enabled in part by support provided by the British Columbia DRI Group and the Digital Research Alliance of Canada (alliancecan.ca). The authors affirm that there are no conflicts of interest to disclose.

## Extended Data

**Extended Data Table 1-1.**
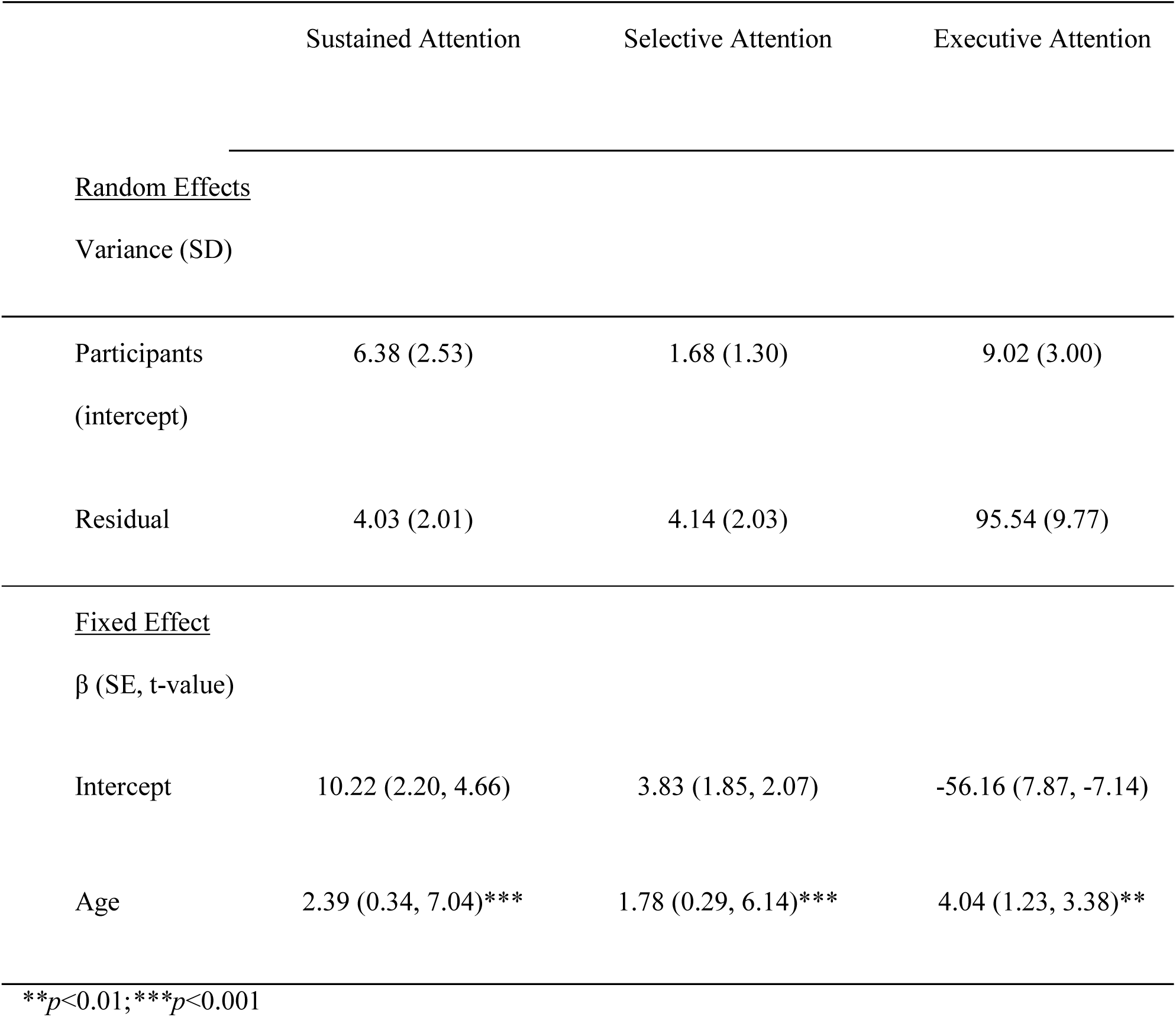
Linear mixed-effects models between age and attention.

**Extended Data Table 2-1.** Parcellation Region Abbreviations and Full Names.

| Region Abbreviation | Region Full Name | Region Abbreviation | Region Full Name |
| --- | --- | --- | --- |
| AntTemp | anterior temporal | PFCd | dorsal prefrontal cortex |
| Aud | auditory | PFCl | lateral prefrontal cortex |
| Cent | central | PFCld | lateral dorsal prefrontal cortex |
| Cinga | cingulate anterior | PFClv | lateral ventral prefrontal cortex |
| Cingm | mid-cingulate | PFCm | medial prefrontal cortex |
| Cingp | cingulate posterior | PFCmp | medial posterior prefrontal cortex |
| ExStr | extrastriate cortex | PFCv | ventral prefrontal cortex |
| ExStrInf | extra-striate inferior | PHC | parahippocampal cortex |
| ExStrSup | extra-striate superior | PostC | post central |
| FEF | frontal eye fields | PrC | precentral |
| FPole | frontal pole | PrCd | precentral dorsal |
| FrMed | frontal medial | PrCv | precentral ventral |
| FrOper | frontal operculum | RSC | retrosplenial cortex |
| IFG | inferior frontal gyrus | Rsp | retrosplenial |
| Ins | insula | S2 | S2 |
| IPL | inferior parietal lobule | SPL | superior parietal lobule |
| IPS | intraparietal sulcus | ST | superior temporal |
| OFC | orbital frontal cortex | Striate | striate cortex |
| ParMed | parietal medial | StriCal | striate calcarine |
| ParOcc | parietal occipital | Temp | temporal |
| ParOper | parietal operculum | TempOcc | temporal occipital |
| pCun | precuneus | TempPar | temporal parietal |
| pCunPCC | precuneus posterior cingulate cortex | TempPole | temporal pole |

**Extended Data Table 2-2.** Cosine similarities of mean-centred task PLS analyses and behavioural PLS analyses with potential confounds.

| Measure | Cosine Similarity with Potential Confounds<br>Behavioural PLS | *p*-value |
| --- | --- | --- |
| SC Local Clustering | -0.11 | 0.47 |
| SC Weighted<br>Degree | 0.25 | 0.13 |
| FC Local Clustering | 0.54 | 0.50 |
| FC Weighted<br>Degree | 0.49 | 0.49 |
| SC-FC Coupling | 0.02 | 0.81 |

The cosine similarity of the brain scores and the *p*-values (based on permutation testing) between each mean-centred task PLS analysis and the behavioural PLS analysis of the respective brain metric with sex and motion metrics. Abbreviations: SC = structural connectivity; FC = functional connectivity; LV = latent variable.

**Extended Data Table 4-1.** Cosine similarities of structural connectivity local clustering behavioural PLS analyses.

| Measure | Cosine<br>Similarity with<br>Task PLS | *p*-<br>value | Cosine Similarity<br>with Potential<br>Confounds<br>Behavioural PLS | *p*-<br>value |
| --- | --- | --- | --- | --- |
| SC Local Clustering -<br>Sustained Attention<br>LV2 | 0.06 | 0.61 | 0.11 | 0.70 |
| SC Local Clustering -<br>Selective Attention | -0.07 | 0.65 | 0.20 | 0.51 |
| SC Local Clustering -<br>Executive Attention | -0.009 | 0.93 | 0.11 | 0.73 |

The cosine similarity of the brain scores and the *p*-values (based on permutation testing) between each behavioural PLS (bPLS) analyses of the structural connectivity (SC) local clustering metric with a) the SC local clustering mean-centred task PLS analysis and b) the bPLS analysis of SC local clustering with sex and motion metrics. Abbreviations: SC = structural connectivity; LV = latent variable.

**Extended Data Table 5-1.** Cosine similarities of structural connectivity weighted degree behavioural PLS analyses.

| Measure | Cosine<br>Similarity with<br>Task PLS | *p*-<br>value | Cosine Similarity with<br>Potential Confounds<br>Behavioural PLS | *p*-<br>value |
| --- | --- | --- | --- | --- |
| SC Weighted Degree<br>- Sustained Attention<br>LV1 | -0.03 | 0.84 | -0.15 | 0.53 |
| SC Weighted Degree<br>- Sustained Attention<br>LV2 | 0.09 | 0.49 | 0.32 | 0.074 |
| SC Weighted Degree<br>- Selective Attention | -0.07 | 0.60 | -0.038 | 0.85 |
| SC Weighted Degree<br>- Executive Attention | -0.02 | 0.89 | 0.023 | 0.91 |

The cosine similarity of the brain scores and the *p*-values (based on permutation testing) between each behavioural PLS (bPLS) analyses of the structural connectivity (SC) weighted degree metric with a) the SC weighted degree mean-centred task PLS analysis and b) the bPLS analysis of SC weighted degree with sex and motion metrics. Abbreviations: SC = structural connectivity; LV = latent variable.

**Extended Data Table 6-1.** Cosine similarities of SC-FC coupling behavioural PLS analyses.

| Measure | Cosine<br>Similarity with<br>Task PLS | *p*-<br>value | Cosine Similarity with<br>Potential Confounds<br>Behavioural PLS | *p*-<br>value |
| --- | --- | --- | --- | --- |
| SC-FC Coupling -<br>Sustained Attention | 0.08 | 0.51 | 0.063 | 0.74 |
| SC-FC Coupling -<br>Selective Attention | 0.07 | 0.83 | 0.63 | 0.031 |
| SC-FC Coupling -<br>Executive Attention | 0.06 | 0.77 | -0.30 | 0.21 |
| SC-FC Coupling, SC<br>Weighted Degree, FC<br>Weighted Degree –<br>Age | NA | NA | 0.14 | 0.35 |

The cosine similarity of the brain scores and the *p*-values (based on permutation testing) between each behavioural PLS (bPLS) analyses with a) the SC-FC coupling mean-centred task PLS analysis and b) the bPLS analysis of SC-FC coupling with sex and motion metrics. Abbreviations: SC = structural connectivity; FC = functional connectivity.

**Extended Data Table 6-2.** Variance explained by the principal components analyses of structural and functional connectivity.

| Principal Component | Structural Connectivity<br>Percentage of Variance Explained | Functional Connectivity<br>Percentage of Variance Explained |
| --- | --- | --- |
| 1 | 89.06 | 67.50 |
| 2 | 0.45 | 1.74 |
| 3 | 0.36 | 1.14 |
| 4 | 0.34 | 1.04 |
| 5 | 0.32 | 0.94 |
| 6 | 0.30 | 0.87 |
| 7 | 0.29 | 0.80 |
| 8 | 0.29 | 0.78 |
| 9 | 0.28 | 0.73 |
| 10 | 0.26 | 0.72 |

The percentage of variance explained by the first ten principal components of the principal components analyses with subjects' structural and functional connectivity matrices.

## Notes

https://github.com/McIntosh-Lab/Rokos2024_SCFC_NetworkAnalyses

### Competing Interest Statement

The authors have declared no competing interest.

### Summary of Updates

Please see the updated revision correctly entering the corresponding author as "Anthony R. McIntosh".

